# Late ERP amplitude for self-face perception positively associated with heartbeat perception accuracy

**DOI:** 10.1101/792366

**Authors:** A.V. Tumialis, T.A. Alikovskaia, A.S. Smirnov, P.P. Khoroshikh, K.A. Fadeev, S.A. Gutnikov, K.S. Golokhvast

## Abstract

Perception of yourself involves the integration of information from various sources. In a number of studies, it was found that the perception of one’s own face is accompanied by an increase in the accuracy of perception of heartbeats and the amplitude of brain potentials caused by heart beats. In this study, subjects had to do a heartbeat count test to determine the accuracy of the interception. Then, the subjects were presented with the faces of an unknown person, a friend and the subject’s own face. The simultaneous registration of EEG was organized. We analyzed the relationship between the amplitude of the evoked potentials when viewing these faces and the accuracy of interoception. It was found that the amplitude of the late EP component (850 - 1106 ms) has a positive correlation with IAcc in the central and right parietal and occipital areas when perceiving one’s own face. According to the localization of distributed sources of activity, it was found that the connection is localized in the right anterior upper temporal cortex. Thus, the association between exteroceptive perception of one’s own face and IAcc occurs in the late period of EP. Moreover it is localized in the right temporal region of the cortex, associated with multisensory integration and recognition of personal information.

## INTRODUCTION

Numerous studies have shown priority in recognition one’s own face compared to faces of unknown people (Keyes et al., 2010; Tacikowski et al., 2011; Wang et al., 2011; Geng et al., 2012). Another important component of identity is the perception of one’s own body (Craig, 2002). However, the specific time of sensory integration between the perception of one’s own face and body remains unknown.

Electrophysiological research found that structural analysis of the face is associated with N170 ERP component (Eimer, 2000). Self-face recognition produce more negative amplitude of this component (Caharel et al., 2005; Keyes et al., 2010; Geng et al., 2012), but some study did not found the same effect (Sui et al., 2006; Tanaka et al., 2006; Wang et al., 2011).

The medial frontal negative amplitude of component N200 reflects the activation of cognitive control mechanisms in the form of action monitoring and error detection (Ullsperger et al., 2014). In line with this view, the activation of the medial frontal cortex corresponds to a mismatch between the image of the self and the unknown person (Flagan & Beer, 2013) reflected in less negative amplitude for the self-face compare to strangers (Keyes et al., 2010;…) and highly important self-related content (Xu et al., 2017).

N250 increases with the perception of one’s own face and is associated with a high degree of representation in the memory of a person’s face (Tanaka et al., 2006) and taps into the long-term perceptual memory for any type (face or non-face) of individuated stimulus (Pierce et al., 2011).

The late components P300 and LPP reflect personal significance (Xu et al., 2017) and are more positive in the perception of one’s own face (Keyes et al., 2010; Tacikowski et al., 2011; Geng et al., 2012).

Tomographic studies showed that in addition to the initial structural analysis of the face wich is associated with activation located in the occipital and posterior inferior temporal cortex (Zhao et al., 2017; Wang et al., 2017), a Second-order analysis of the face identity in ATL is taking place.

Activation of this cortical area occurs independently of rotation (Anzellotti et al., 2014) and half of the perceived face (Anzellotti et al., 2016) and reflects association between face and person identity representations (Wang et al., 2017) and person-related semantic information in memory (Tsukiura et al., 2008). The results demonstrate that the recognition of an individual face causes activation on the right, and the connection of the person with one’s name activates ATL on the left (Pobric et al., 2016). Moreover, the dorsal and ventral ATL roles are dissociable between two steps of association, associations of person-related semantics with name and with face (Tsukiura et al., 2010). ATL also plays an important role in higher-level information processing and is associated with social cognition (Olson et al., 2013) and stable abstract social conceptual representations or moral norms (Zahn et al., 2009). Activation in right ATL in the perception of one’s own face was found in studies by Uddin et al., (2005) and Platek et al., (2006).

Also recognition (Denny et al., 2012) and involuntary self-assessment (Egenolf et al., 2013) involve activation of medial prefrontal (MPF) and anterior cingulate (ACC) cortices as parts of the Default Mode Network (Davey et al., 2016). Moreover, the activation of latter areas of the cerebral cortex is also manifested not only by the perception of the images or verbal prompts, but also accompanied by the appearance of spontaneous thoughts about oneself (Knyazev, 2013).

Never the less, the temporal and spatial characteristics of the brain reaction in response to the presentation of the subject’s face leave open the question of the place and time of communication with the perception of one’s own body.

Interception is often operationalized as the accuracy of the perception of heartbeats (Schandry, 1981). In a number of studies, IAcc is seen as upward activation with localization in the right anterior superior insular cortex (Craig, 2002; Critchley et al., 2004). This approach is supported by evidence of the association of IAcc and amplitude of Heartbeat Evoked Potential (Pollatos et al., 2005). However, this view is criticized (Brener, J., & Ring 2016) because false feedback increases the accuracy of interception (Ring et al., 2015) and causes activation of the same areas of the cortex (Kleint et al., 2015) as the real one perception of heartbeats.

An alternative approach is being developed as part of the predictive coding approach (Seth 2013, Barrett, L. F., & Simmons 2015; Ainley et al., 2016), in which the IAcc is considered as comparing the subject’s predictive model of his own body with the data of upward activation of the autonomic nervous system.

Despite the conceptual disagreements in the approaches, positive associations of IAcc with the amplitude of the late EP components were found in studies of the perception of emotional images (Herbert et al., 2007) and attention (Pollatos et al., 2007). Positive correlations of IAcc with the amplitude of the late positive component of the error response were also found (Sueyoshi et al., 2014, Godefroid et al., 2016). In a study by Fischer et al., (2013), subjects performed the Go-NoGo task from a first-person perspective and from the perspective of another person. It was found that making mistakes from a third-person perspective did not lead to the appearance of a brain potential caused by an error. If an error is nevertheless made and attributed to itself, the formation of a late positive reaction complex to the error is formed.

In the study of the connection between the perception of one’s own face and body signals, significant effects were also found. In particular, subjects with higher IAcc have a more negative amplitude of the early HEP component when perceiving a face containing 40% of the subject’s own face (Sel et al., 2017). They are also less susceptible to the illusion of owning a face (Tsakiris et al., 2011). Therefore, in subjects with high IAcc, it increases sensitivity to the signals of one’s own body during the perception of signs of one’s own face and decreases the possibility of identification with the face of another person.

Another series of studies showed that self-perception increases IAcc regardless of the emphasis on the bodily or autobiographical Self (Ainley et al., 2013), but does not change in East Asians (Maister et al., 2014) and is reduced in those who suffering from anorexia nervosa (Pollatos et al., 2016).

Thus, the perception of one’s own face increases sensitivity to heartbeats, and this connection probably occurs in the late stages of the brain’s response to exteroceptive stimuli. At the same time, we did not find studies of the relationship between IAcc and the amplitude of the EP in the perception of the subject’s own face.

In this study, we evaluated the magnitude of the accuracy of the interception of subjects using the widely used test of accuracy of interception in the variant of counting heart beats (Schandry 1981). Next, subjects were presented with photographs of faces of an unknown person, friend and face of the subject himself, and an electrophysiological reaction of the brain was recorded in parallel. We suggest that the early components are associated with perceptual identification mechanisms, while the later ones may be associated with the accuracy of interception, since it requires the activation of a body model. We also assume that the correlation will be higher upon presentation of one’s own face, as bodily identity is activated. When perceiving the face of a friend and an unknown person, the correlation will be lower since they do not activate the subject’s bodily identity.

## METHOD

### Participants

Thirty subjects took part in the study (age: M=19.17 SD=1.28; males: N = 11; education: M=12.28, SD=1.31; BMI: M=21.43, SD=2.87). Subjects refrained from taking alcohol for 1 day and tobacco and caffeine 2 hours before the study. Participation was voluntary and all subjects signed informed consent. The study was carried out in accordance with the declaration of Helsinki and approved by the FEFU bioethical committee.

### Interoceptive accuracy

The heartbeat perception task was performed according to the mental tracking method by Schandry (1981) using three intervals of 25, 45, 60 s that were separated by periods of 10 s. Increasing and decreasing sequences were excluded and the remaining sequences variants were randomised between subjects. Participants were asked do not move during the registration, keep their eyes open and count their own heartbeats silently. The beginning and the end of the counting intervals were signalled acoustically. After the stop signals participants were asked to verbally report the number of counted heartbeats. The participants were not informed about the length of the counting phase nor about the quality of their performance. Participants were not permitted to take their pulse or to attempt any other manipulations that could facilitate the detection of heartbeats.

Interoceptive accuracy (IAcc) was estimated as the mean heartbeat perception score according to the following transformation:

n – number of periods, RH - Recorded Heartbeats, CH - Counted Heartbeats.

Maximum is equal 1 and correspond to absolute accuracy of heartbeat perception.

### Stimuli

Each subject sent photos of his or her own face and the photo of his or her friend’s face of the same sex. The faces on the photoes should be emotionless and should be clearly identified (without gross shadows or hairs covering the face). The faces from Karolinska Directed Emotional Faces (KDEF) with neutral expression, four females (AF01NES, AF11NES, AF19NES, AF34NES) and four males (AM10NES, AM17NES, AM23NES, AM26NES), were taken as control condition. All photos were cut off on the upper edge of the hairstyle and the lower part of the neck. Finally, the photos were discolored and aligned in contrast and brightness. The size of the images was 23 × 16 cm, which from a distance of 1 meter was 13°7’x 9°9’. All photos were shown against a black background.

### Procedure

Registration took place in the room with controlled lighting and each participants was sitting on a chair at a distance of 1 meter in front of the monitor. Four faces of unknown persons of the same sex as the participants were given to the subject with the instructed to choose one that does not cause positive or negative emotional associations and does not attract attention and will not be remembered by chance meeting. The choice of one person is due to the fact that the participant’s and the friend’s photo were also in the singular. Then the instruction followed, during which all three faces (the participant’s own face, participant’s friend face and the face of unknown person) after appropriate processing were presented to the participants of the experiment were presented and asked them to watch these faces during the task.

In the course of the experiment the faces were presented in the random order, each face been presented 32 times. The sequence of the events in trial was as follows: a white cross in the center of the screen for 1 second, then a face presentation for 5 seconds and an interstimulus interval for 1.5 seconds. Paradigm programming and stimuli presentation were carried out in PsychoPy 1.82.

Photosensor in the upper left corner of the screen was used for precise synchronization of stimuli presentation with the EEG.

### EEG

EEG acquisition was performed using an NVX-52 (https://mks.ru/en/products/nvx/) amplifier using 44 point Ag/AgCl electrodes with a ground at AFz. The sampling rate was 1 kHz. Electrode resistance was below 10 kOhms for all electrodes. The EEG was recorded against the averaged values of the ear lobes reference electrodes. Online high-pass filter was 0.05 Hz and a low-pass filter was 100 Hz.

Horizontal eye movements were recorded with electrodes placed at the outer canthus of each eye. Vertical eye movements were recorded with electrodes placed above and below left eye. The band-pass filter for EOG recording was 1-20 Hz.

ECG was recorded using the two external channels with a bipolar ECG lead II configuration. ECG data were offline band-pass filtered between 1 and 40 Hz.

### Event Related Potentials

For the ERP analysis, the continuous EEG data were filtered between 0.1 and 15 Hz, rereferenced to common average and epoched from −200 to 1500 ms from the beginning of the stimulus presentation. Segments were baseline corrected for a pre-stimulus interval of −200 to 0 ms. Epochs were visually inspected and in the case of presence of pronounced artifacts were removed. Next, the individual components were extracted using the Infomax algorithm. The removal of the components containing EOG and myographic artifacts was carried out semi-automatically. Further, the epochs were again visually inspected and, exceeding the threshold of 80 mV, were removed. Finally, EEG data were averaged for each category.

The analysis was carried out in T5/T6 for the N170 in the period from 140 to 200 ms, in the Cz electrode for N200 from 200 to 320 ms, in PO7/PO8 for the N250 in the period from 230 to 330 ms. The analysis of the late components was carried out in the CPz electrode for P300 in the period 400 – 600 ms and for LPP in the period 600 – 1000 ms.

Further data were analysed using the Fieldtrip software package (http://www.ru.nl/fcdonders/fieldtrip/), a Matlab-based toolbox for the analysis of electrophysiological data. The differences between the groups were estimated using cluster-based permutation statistics with the data driven approach. (These statistics are based on clustering of adjacent time-electrode samples that all exhibit a similar difference in sign and magnitude.) It is a nonparametric test and does not require assumptions about the normality of data distribution.

The procedure was as follows. First, t-test differences (p < 0.05) were calculated between the groups in each electrode and at each time point in the period from −0.05 to 0.6 seconds relative to the timing of the motor response. Based on spatiotemporal adjacency (minimum of 2 neighbouring units) clusters were created and a cluster with a maximum sum of t-values was used in further statistics. Next, the CSD data was permuted between the groups and the statistics were calculated again. A total of 2000 randomizations was used and a permutation distribution of the maximum differences in the clusters was created. Further, the probability of deviating the data obtained in the study from the parameters of this distribution using the Monte Carlo method was done to calculate the significance level (p < 0.025). This approach allows us to exclude the acceptance of false positive results in multiple comparisons and to allocate clusters by time and space, based on the experimental data.

### Source localization

Based on the distribution of electric potential recorded on the scalp, the standardized low resolution brain electromagnetic tomography (sLORETA) software (publicly available free academic software at http://www.uzh.ch/keyinst/loreta.htm) was used to compute the cortical three-dimensional distribution of current density. sLORETA uses a distributed source localization algorithm to solve the inverse problem of brain electric activity regardless of the final number of neural generators (Pascual-Marqui et al., 2002). The sLORETA algorithm calculates the current density values (unit: amperes per square meter; A/m^2^) of 6.239 gray matter voxels belonging to the brain compartment with a spatial resolution of 5 mm × 5 mm × 5 mm each. The whole three-dimensional brain compartment comprises cortical gray matter and the hippocampus only and does not contain any deep brain structures such as the thalamus or the cerebellum. In the present study, the source location was investigated using voxel-wise randomization tests with 5000 permutations based on statistical non-parametric mapping. To identify possible differences in neural activation between the face categiries non-parametric statistical analyses of sLORETA images (Statistical non-Parametric Mapping; SnPM, Nichols and Holmes, 2002) was performed while employing a *t* statistic (on log-transformed data). The results corresponded to maps of *t* statistics for each voxel, for the corrected *p* < 0.05.

### Statistics

Analysis of the event related potentials was carried out with ANOVA included the Face (Other, Friend, Self) as within subject factor. Sphericity was controlled with the help of the Greenhouse-Geisser criterion, the magnitude of the effect was estimated using eta partial in the square, and post-hoc differences were calculated using the Bonferroni test. Analysis of the correlations was performed using the Pearson r coefficient. All variables were checked for normal distribution with the Kolmogorov-Smirnov test. The significance level was p < 0.05.

## RESULTS

### Heartbeat perception accuracy

HBP scores (Mean = 0.686, SD = 0.178, Range = 0.366 – 0.973, CI = 0.620-0.753) did not differ from normal distribution (K-S d = 0.114, p > 0.20). Also HR data (Mean = 75.15, SD = 10.89, CI = 71.11 – 79.26) did not differ from normal distribution (K-S d = 0.180, p > 0.20). Correlation between HBP and HR was negative (r = −0.42, p = 0.021).

### ERP

First, we present the traditional statistical analysis of ERP amplitude data. Mean, standard deviation and confidence interval data presented in Table 1 and ERP waveforms in Figure 1.

**Table 1.**
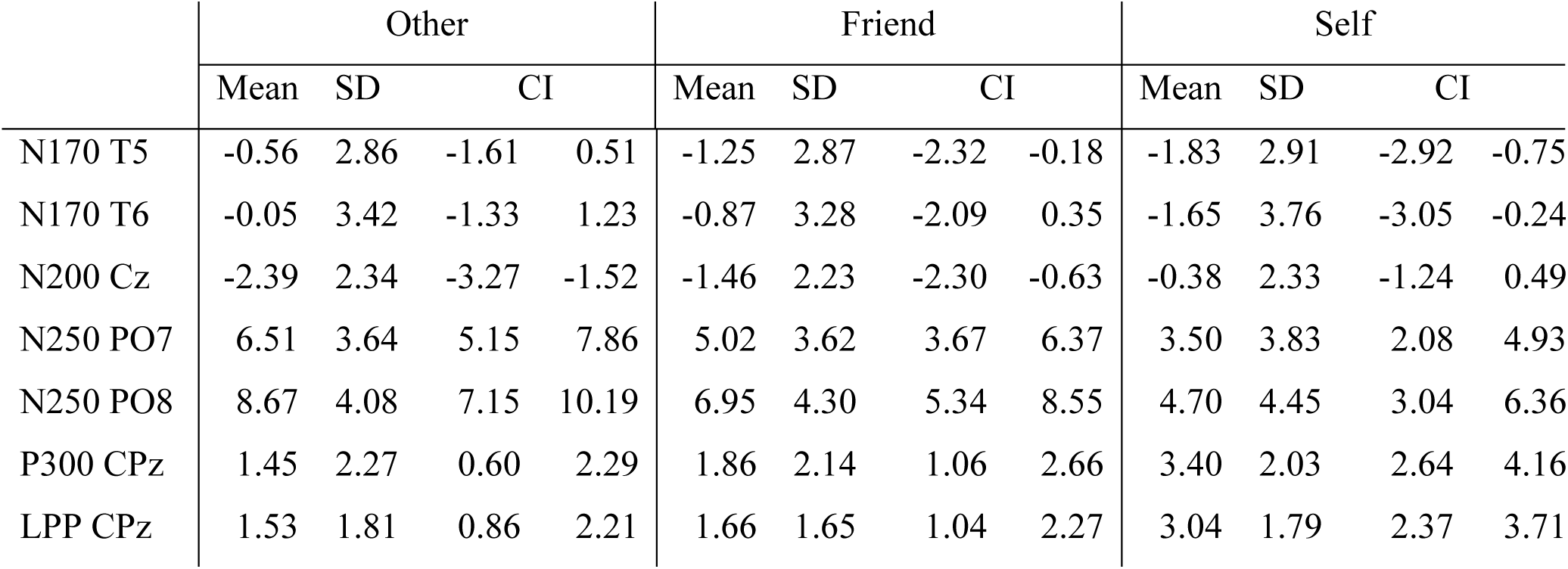
Amplitude statistics of ERP components, evoked by the Other, Friend and Self faces.

|  | Other |  |  |  | Friend |  |  |  | Self |  |  |  |
| --- | --- | --- | --- | --- | --- | --- | --- | --- | --- | --- | --- | --- |
|  | Mean | SD | CI |  | Mean | SD | CI |  | Mean | SD | CI |  |
| N170 T5 | -0.56 | 2.86 | -1.61 | 0.51 | -1.25 | 2.87 | -2.32 | -0.18 | -1.83 | 2.91 | -2.92 | -0.75 |
| N170 T6 | -0.05 | 3.42 | -1.33 | 1.23 | -0.87 | 3.28 | -2.09 | 0.35 | -1.65 | 3.76 | -3.05 | -0.24 |
| N200 Cz | -2.39 | 2.34 | -3.27 | -1.52 | -1.46 | 2.23 | -2.30 | -0.63 | -0.38 | 2.33 | -1.24 | 0.49 |
| N250 PO7 | 6.51 | 3.64 | 5.15 | 7.86 | 5.02 | 3.62 | 3.67 | 6.37 | 3.50 | 3.83 | 2.08 | 4.93 |
| N250 PO8 | 8.67 | 4.08 | 7.15 | 10.19 | 6.95 | 4.30 | 5.34 | 8.55 | 4.70 | 4.45 | 3.04 | 6.36 |
| P300 CPz | 1.45 | 2.27 | 0.60 | 2.29 | 1.86 | 2.14 | 1.06 | 2.66 | 3.40 | 2.03 | 2.64 | 4.16 |
| LPP CPz | 1.53 | 1.81 | 0.86 | 2.21 | 1.66 | 1.65 | 1.04 | 2.27 | 3.04 | 1.79 | 2.37 | 3.71 |

**Figure 1.**
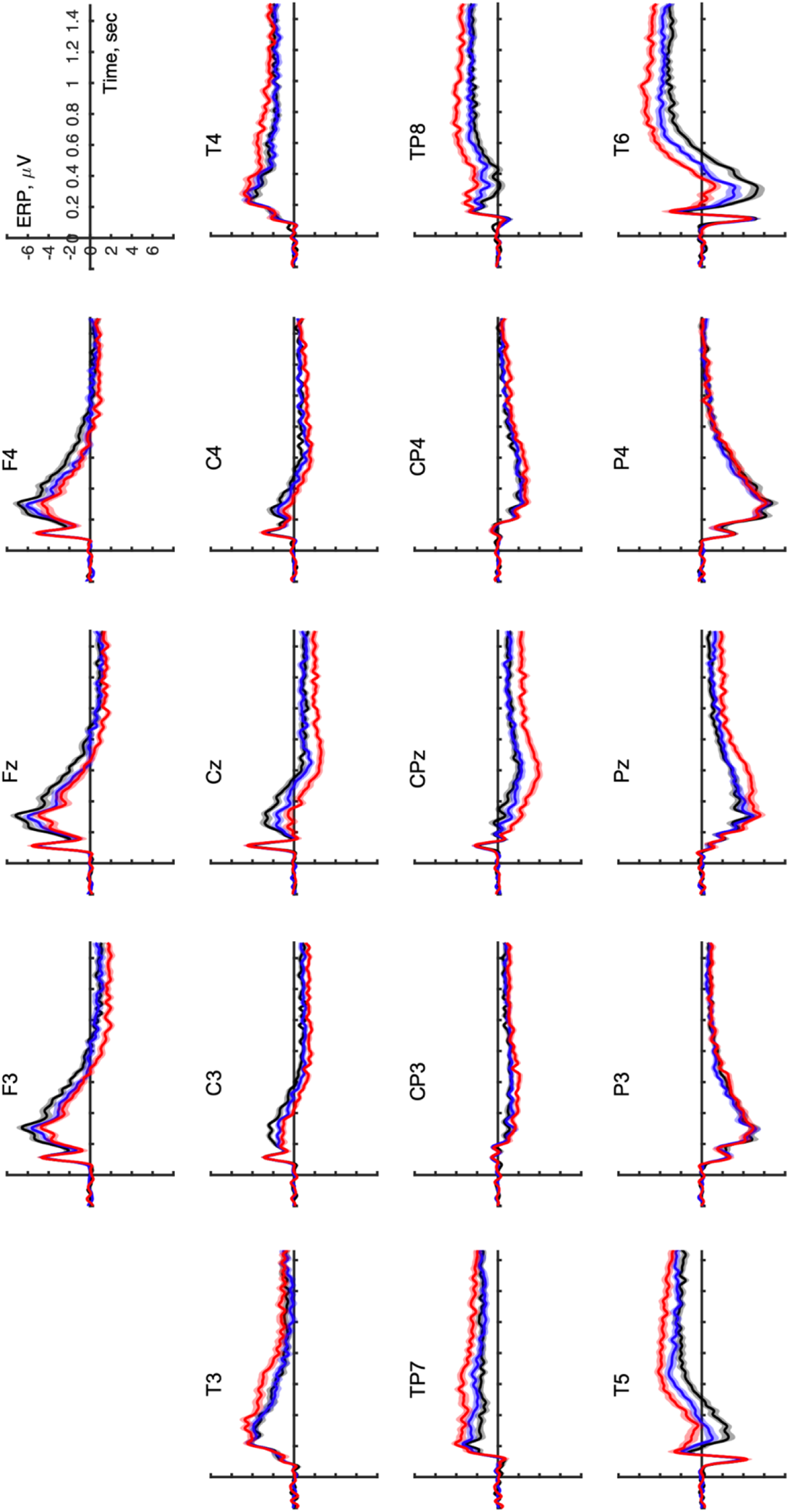
ERP waveforms for Other (black line), Friend (blue line) and Self (red line) faces. Translucent areas depict standard error of mean.

### N170

The face effect was found (F (2.58) = 16.45, p <0.001, Gg = 0.835, Eta = 0.362). The effects of the hemisphere (F (1.29) = 0.25, p = 0.621) and the interaction of the hemisphere and the face (F (2.58) = 0.27, p = 0.762) did not reach a significant level. Post-hoc comparisons showed that the face of an unknown person causes the greatest amplitude, a smaller amplitude for the friend’s face and the smallest for the subject’s own face were found. All differences ps <0.03 (Bonferroni corrected).

N200 showed a significant face effect (F (2, 58) = 30.97, p <0.001, Gg = 0.950, Eta = 0.516), post-hoc comparisons showed that the face of an unknown person causes a negative amplitude, the face of a friend is smaller and the amplitude for the subject’s face was near zero, all p <0.005.

N250 has significant face effects (F (2, 58) = 52.10, p <0.001, Gg = 0.855, Eta = 0.642), hemisphere (F (1, 29) = 7.16, p = 0.012, Gg = 1.000, Eta = 0.198) and the interaction of the face and hemisphere (F (2, 58) = 3.57, p = 0.038, Gg = 0.938, Eta = 0.110). The most positive values of the amplitude were found in the perception of an unknown person, smaller in the perception of the face of a friend and smallest in the perception of the subject’s own face, all p <0.001. On the right, the amplitude was more positive than on the left. Comparison of the average values of the amplitudes between the hemispheres during the perception of each face revealed that the amplitude of N250 is greater on the right than on the left when perceiving an unknown face (t = - 2.90, p = 0.007) and the face of a friend (t = −2.93, p = 0.007) and the amplitude is not differ between hemispheres in the perception of their own faces (t = −1.78, p = 0.086).

P300 showed a significant face effect (F (2, 58) = 25.17, p <0.001, Gg = 0.966, Eta = 0.465). Post-hoc comparisons revealed that one’s own face causes a more positive amplitude compared to faces of an unknown person and a friend, ps <0.001.

The LPP amplitude also showed a face effect (F (2, 58) = 12.69, p <0.001, Gg = 0.964, Eta = 0.304). Own face causes a more positive amplitude compared to faces of an unknown person and a friend, ps <0.001.

Thus, N170 and N250 decreases sequentially, and N200 increases from the face of an unknown person through the face of a friend to the subjects’ own faces. Later components P300 and LPP showed the same result - own face causes a more positive amplitude compared to both other faces.

Because EEG analysis with traditional statistical methods is susceptible to the acceptance of a large number of false positives, known as the multiple comparisons problem, so, group difference would be a false effect. To resolve this problem, we used more robust cluster-based permutation statistics for investigating the differences between ERP amplitudes for the faces. This significantly reduces the acceptance of false effects, since it is distributed on the basis of the maximum sums of t-differences within the space-time clusters in each iteration of the permutations.

Testing the ERP amplitude in the latency range from 0 to 1500 ms the cluster-based permutation test was performed between the Other vs Friend, Other vs Self and Friend vs Self face conditions. We found three negative clusters for Self vs Other, two negative clusters for Friend vs Other and one positive cluster Self vs Friend differences (p < 0.025). See table 2

**Table 2.** Cluster-based permutation statistical differences between ERP amplitudes, evoked by the Other, Friend and Self faces.

|  |  | Time, ms | p | clusterstat | SD | CI range |
| --- | --- | --- | --- | --- | --- | --- |
| Self vs Other | CI 1 | 202 - 610 | 0.0005 | -15903.0778 | 0.0005 | 0.0010 |
|  | CI 2 | 1240 - 1373 | 0.0335 | -1644.8505 | 0.0040 | 0.0079 |
|  | CI 3 | 762 - 804 | 0.0480 | -1316.0925 | 0.0048 | 0.0094 |
| Friend vs Other | CI 1 | 279 - 725 | 0.0005 | -13050.9174 | 0.0005 | 0.0010 |
|  | CI 2 | 148 - 265 | 0.0155 | -2890.0061 | 0.0028 | 0.0054 |
| Self vs Friend | CI 1 | 556 - 743 | 0.0100 | 2778.4055 | 0.0022 | 0.0044 |

The results indicate that ERP amplitude for Friend and Self faces were less positive over the occipital and lateral parietal electrodes compare with Other face from 200 to more than 700 ms and for the Self-face in a later LPP period. Also ERP amplitude for Self-face was more positive then Friend-face over the parietal area in LPP time range (Figure 2).

**Figure 2.**
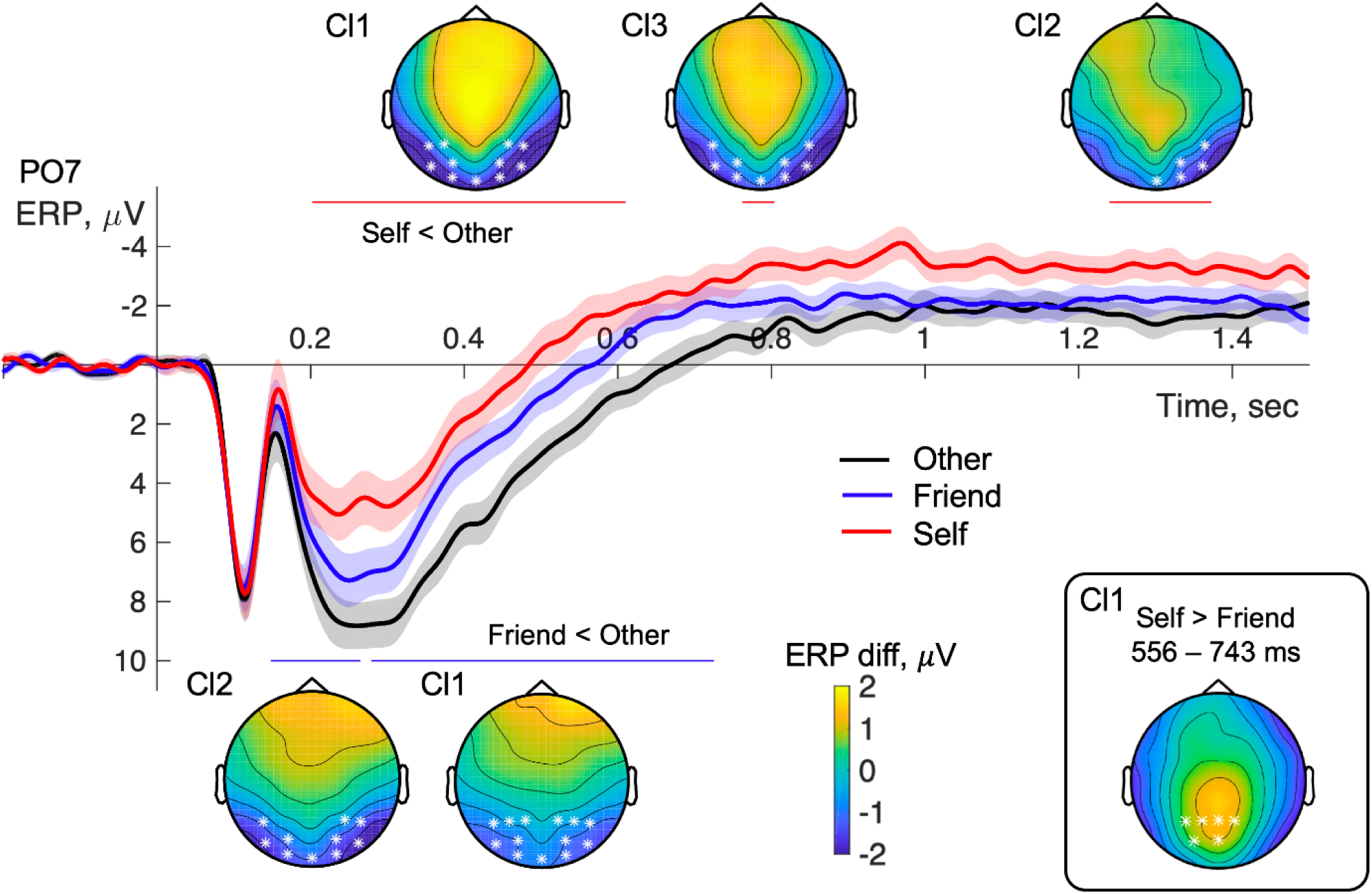
Cluster topoplots for statistical differences between ERP amplitudes evoked by the Other, Friend and Self faces. Asterisks indicated clusters of differences (p < 0.025). Red lines above waveform show time periods of clusters of differences between Self and Other faces. Blue lines below waveform show time periods of clusters of differences between Friend and Other faces. Insertion in bottom right corner show the cluster of differences between Self and Friend faces. Translucent areas around waveforms depict standard error of mean.

### Correlations between ERP and HBP

The cluster-based permutation correlation test was performed between HBP scores and ERP amplitude for Self-face condition in the latency range from 0 to 1500 ms. We found two negative and five positive clusters and only one had a p-value less than 0.025. Within this cluster (clusterstat = 4473.36, Monte-Carlo p = 0.0115, SD = 0.0024, CI range = 0.0047) the positive correlation was significant in the latency range from 850 to 1106 ms over the parietal and occipital electrodes, lateralized to the right (Figure 3). The results indicate that subjects with higher HBP scores have a greater ERP amplitude for self-face perception than subjects with lower HBP scores.

**Figure 3.**
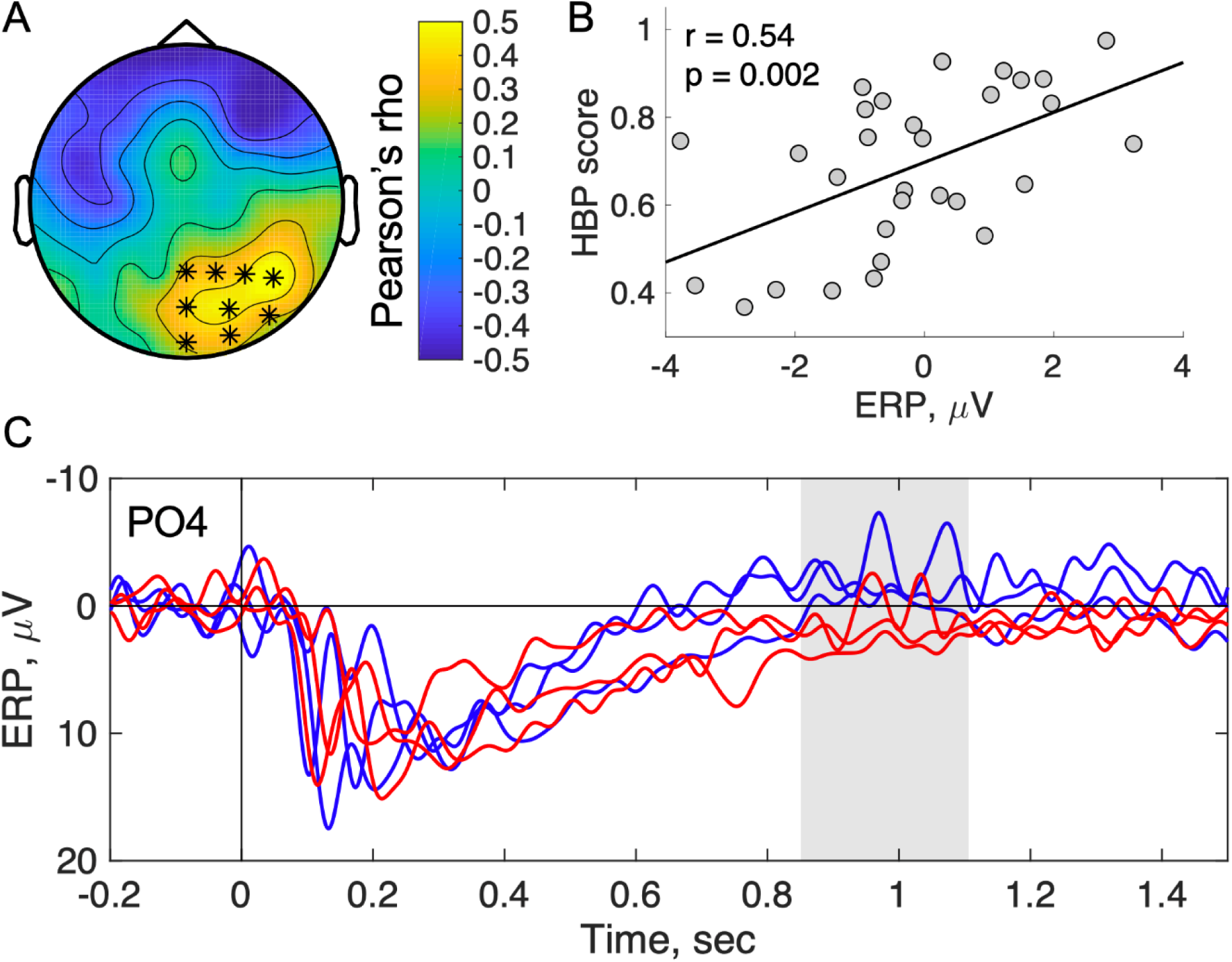
Correlations between ERP amplitude and HBP scores. A. Topographic map of cluster of significant correlations (p < 0.025). B. Scatterplot of correlation between mean amplitude in space and time within the cluster and HBP score. C. Waveforms of three subjects with highest HBP scores (red lines) and lowest HBP scores (blue lines). Gray area indicate the time period of significant cluster correlations.

**Figure 4.**
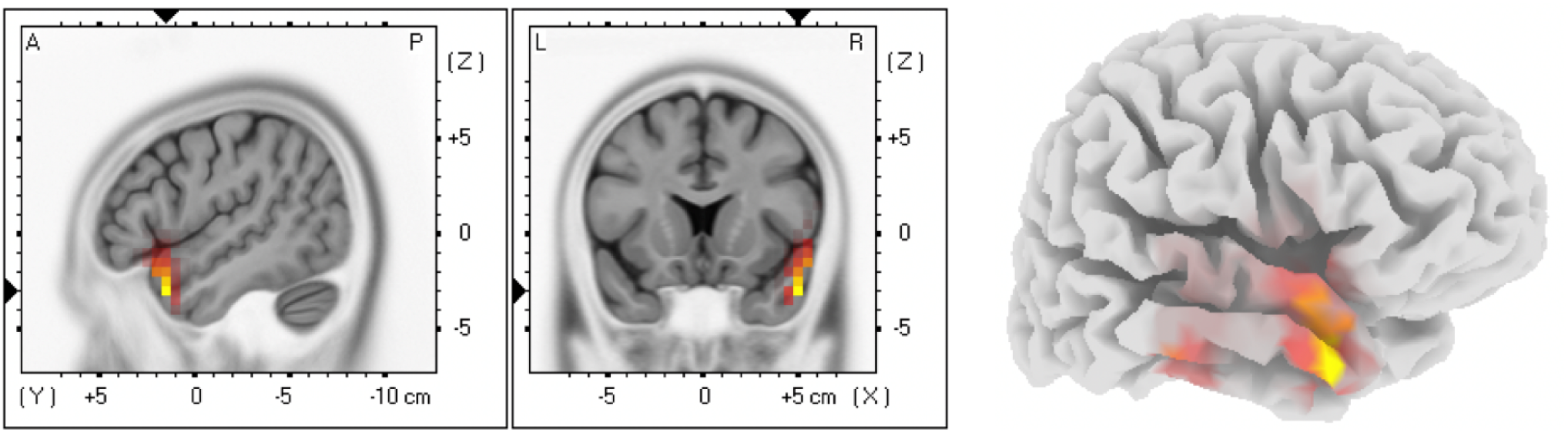
sLORETA statistical nonparametric map correlating the log current source density and HBP scores. Maximum correlation was found at right anterior superior temporal gyrus.

Similar analysis, performed on Other and Friend faces did not reveal significant clusters.

Since HBP scores has a significant correlation with HR, we analyzed the relationship between ERP and HR and found a weak correlation (r = -0.23, p = 0.215).

### Regression analysis

Additional correlation analysis was performed between HBP scores, ERP amplitude within significant cluster from one hand and anxiety, depression and emotional intelligence scores from another hand. Results presented in table 3.

**Table 3.**
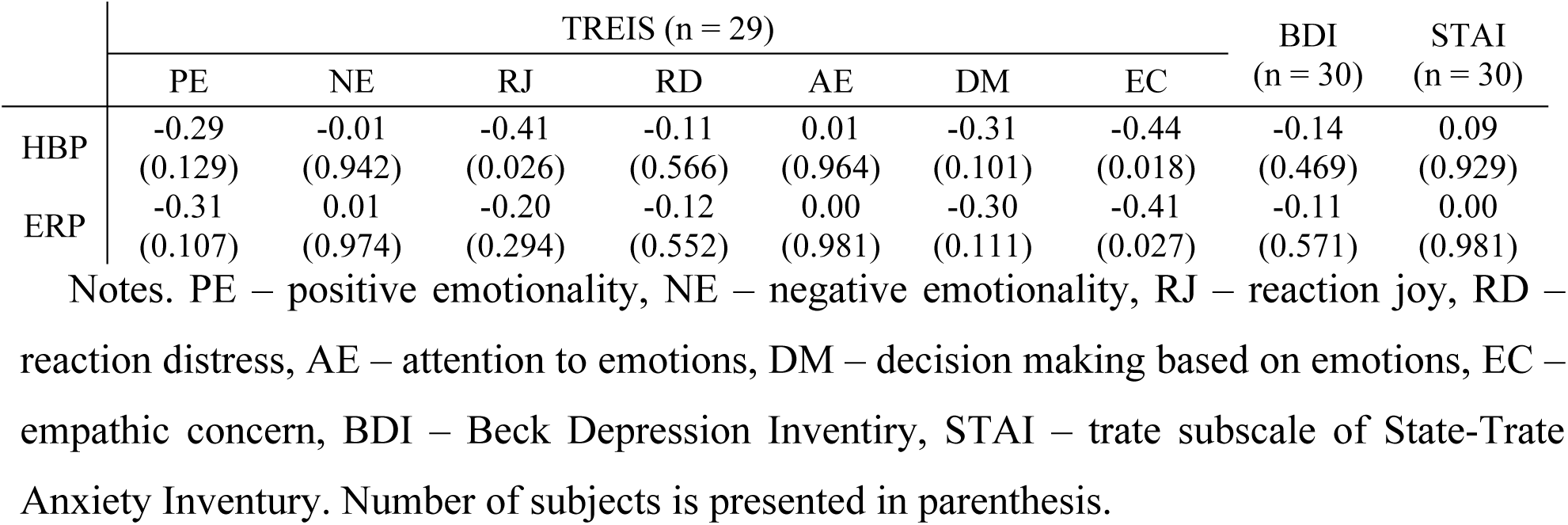
Correlations (Pearson product-moment correlation coefficients) between heartbeat perception (HBP) scores, within-cluster mean ERP amplitude for the Self-face perception and psychometric scales (p-value in parenthesis).

Both HBP scores and ERP amplitude were correlated with the Empathic Concern subscale of TREIS. Thus, regression analysis was conducted to predict ERP amplitude by the HBP scores, Empathic Concern scores and Heartrate. Variables did not differ from normal distribution (K-S test; ERP: d = 0.087; EC score: d = 0.063; all ps > 0.2).

Regression results reveal that ERP amplitude was predicted (Multiple R = 0.58, Multiple R^2^ = 0.34, Adjusted R^2^ = 0.26, F(3,25) = 4.31, p = 0.014) by the HBP scores only (Table 4). Variance inflation factor was 1.44 for HBP scores, 1.24 for EC scores and 1.22 for HR indicating low collinearity. Residuals was normally distributed (K-S test, d = 0.118 p > 0.2).

**Table 4.** Results of multiple regression analysis. Mean within-cluster ERP amplitude for Self-face was predicted variable and heartbeat perception (HBP) scores, Empathic Concern (EC) scores and heartrate (HR) were predictors.

| | b | SE | t | p | CI | | $\beta$ | SE | CI | | Partial R |
| --- | --- | --- | --- | --- | --- | --- | --- | --- | --- | --- | --- |
| intercept | -1.68 | 3.42 | -0.49 | 0.627 | -8.73 | 5.36 |  |  |  |  |  |
| HBP | 4.48 | 1.87 | 2.40 | 0.024 | 0.63 | 8.32 | 0.47 | 0.19 | 0.07 | 0.87 | 0.43 |
| EC | -0.05 | 0.04 | -1.16 | 0.256 | -0.14 | 0.04 | -0.21 | 0.18 | -0.58 | 0.16 | -0.23 |
| HR | 0.002 | 0.03 | 0.08 | 0.934 | -0.06 | 0.06 | 0.01 | 0.18 | -0.35 | 0.38 | 0.02 |

This result indicate that EC scores and heartrate has low influence on ERP amplitude.

Additional factorial forward regression reveal, that only HBP scores predict ERP amplitude, indicating, that any interactions between variables were non significant predictors.

### Localization of activity

A study of the relationship between HBP and the amplitude of activity sources in the selected time showed the presence of significant correlations in the right anterior upper temporal cortex. The coordinates of the maximum correlation are presented in table 5.

**Table 5.** Coordinates and structure of maximum sLORETA statistical nonparametric correlation between the log current source density and HBP scores.

| MNI |  |  | Talairach |  |  | Value | BA | Structure |
| --- | --- | --- | --- | --- | --- | --- | --- | --- |
| X | Y | Z | X | Y | Z |  |  |  |
| 50 | 15 | -30 | 50 | 13 | -26 | 0.672 | 38 | STG |

## DISCUSSION

In this study, we studied the relationship between brain EP in response to the presentation of the subject’s own face, the face of a friend and an unknown person, and the accuracy of perception of heartbeats. Two main results were found. Firstly, the late component of the evoked potential in response to the presentation of the subject’s own face is positively associated with the accuracy of perception of heartbeats, and secondly, the localization of this connection was found in the upper right anterior temporal cortex.

Analysis of the amplitude of the EP showed the presence of effects of the category of the face from the early components N170 and N200 up to the late positive components of the EP. Own face causes a more negative amplitude of N170 and a more positive amplitude of N200, N250, P300 and LPP compared to the face of an unknown person. The face of a friend causes the EP amplitude intermediate between the own face and the face of an unknown person in the early components N170, N200 and N250 and does not differ from the face of an unknown person in the late components P300 and LPP.

The amplitude of N170 is related to the analysis of the configuration of a first-order face (Eimer, 2000). The more negative amplitudes of this component in the perception of the subject’s own face, found in this study, are consistent with previously obtained data (Caharel et al., 2005; Keyes et al., 2010). In addition, a more negative amplitude of N170 when perceiving the subjects’ own faces was found when repeating representations of faces (Tacikowski et al., 2011) when perceiving a photograph of ones’ self compared to reflection in a mirror (Butler et al., 2012). In a number of other studies, the influence of the face was not detected (Sui et al., 2006; Tanaka et al., 2006; Wang et al., 2011). The significant effect in this study may be related to the size of the stimuli. In this and other studies (Caharel et al., 2005; Keyes et al., 2010), the faces were quite large (13 ° 7’, more than 5 ° and ∼ 14 °). In studies without side effects, the incentives were small: 3 ° x3 ° (Sui et al., 2006), 3 ° x4.8 ° (Tanaka et al., 2006) and 2.9 ° x 2.9 ° (Wang et al.., 2011). This issue requires further study.

The medial frontal negative amplitude of component N200 reflects the activation of cognitive control mechanisms in the form of action monitoring and error detection (Ullsperger et al., 2014). In line with this view, the activation of the medial frontal cortex corresponds to a mismatch between the image of the self and the unknown person (Flagan & Beer, 2013). ERP data reveal less negative amplitude for self-face compared face of unknown person () and present results corresponds to these data.

N250 detects second-order relational information seems to be specific for face (Tanaka et al., 2006) or person-associated things (Pierce et al., 2011).

The later EP components P300 and LPP showed a more positive amplitude in the perception of the own face, which reflects the significance (Xu et al., 2017), more intense (Sui et al., 2006) and sustained (Tacikowski et al., 2011) attention.

Traditional statistical analysis is subject to the influence of errors of the first kind - the adoption of false positive results with a large number of comparisons. To reduce this effect, we undertook a more complicated complicated analysis using cluster-based permutation statistics. The study of ERP amplitude differences revealed clusters of amplitude differences in response to an unknown face compared to the faces of the fried and the subjects’ own faces in the parietal-occipital and parietal-temporal areas of the head surface for a period of about 200 to more than 600 ms. The difference in amplitudes in response to one’s own face and unknown face continues in later clusters up to 1373 ms. Also, differences in amplitudes in response to one’s own face and a friend’ face during the LPP period were found in the middle parietal region.

Thus, the ERP results found in this study reflect the known data and, therefore, the technique is valid for use in the study of the relationship between the accuracy of heart rate perception and ERP amplitude. It was the main goal of this study.

In the tasks of attention (Pollatos et al., 2007), perception of emotional images (Herbert 2007) and reaction to errors (Sueyoshi et al., 2014, Godefroid et al., 2016), the accuracy of perception of heartbeats had positive connections with the late components of brain reactions. The subjective image of the body is based on the perception of signals from within the body. Thus, the accuracy of the perception of heartbeats is associated with an increase in the early negative component of the reaction of the brain to heartbeats (Sel et al., 2017). In the present study, we found a positive relationship between the amplitude of the EP during the perception of the own face and the accuracy of perception of heartbeats in the period 850-1106 ms in the central and right parietal-occipital area.

In light of various approaches to interception, the results can be interpreted in various ways. The upstream activation approach suggests that subjective differences in IAcc reflect differences in sensitivity to body signals. The predictive coding approach involves comparing the predictive model of physiological reactivity with the perception of bodily signals. It is difficult to say in favor of which approach the results of this study indicate, since the design of the study and the correlation nature of the results cannot provide an answer to this question. Subsequent studies with a more suitable design may shed light on this issue.

Perceptual characteristics of the face are recognized in the early stages of perception and are manifested in the effects of ERP components. In the late period, the integration of information from other sensory modalities occurs and interception is one of the sources of confirmation of identity. Thus, the perception of oneself in the photo is multisensory in nature. The reaction to the image of one’s own face is associated with the magnitude of the contribution of bodily sensitivity to personal identity. This conclusion is supported by data on increasing the accuracy of perception of heartbeats when healthy subjects of Western culture perceive themselves (Ainley et al., 2013; Maister et al., 2014; Pollatos et al., 2016). But if behavioral assessments were used in these studies, then in this study the accuracy of interoception is associated with an objective reaction obtained using EEG.

Localization of the relationship between the accuracy of the perception of heartbeats and the amplitude of ERP was found in the right anterior temporal cortex with the maximum connection in the superior gyrus.

The anterior temporal lobe (ATL) play a critical role in representing and retrieving social knowledge (Olson 2013). Also, activation of ATL, to a greater extent on the right (Pobric et al., 2016), is associated with the perception of individual faces (Kriegeskorte et al., 2007; Anzellotti, S., & Caramazza 2016). In addition, right upper temporal cortex activation was found for abstract social conceptual representations (Zahn et al., 2009) and abstract person identity representations (Wang 2017). Thus, the results of this study support the interpretation of interception as a personality concept, which corresponds to the predictive coding approach (Seth 2013, Barrett, L. F., & Simmons 2015; Ainley et al., 2016). However, the data do not preclude the possibility of a connection between interception and upward activation, for example, analyzed using HEP (Sel 2017) with internal attention.

In a study by Anzellotti, S., & Caramazza (2017), multisensory integration between visual perception of the face and auditory perception of speech was found in the posterior superior temporal gyrus. In this study, multisensory integration between visual information and interception is found in the anterior superior temporal cortex. Since the right anterior insular cortex has the closest links with interoception (Craig, 2002; Critchley et al., 2004), it is possible that the functional connection of the anterior temporal cortex with the right anterior insula will underlie the association found.

The study found that the perception of one’s own face includes the actualization of bodily identity. At the same time, a violation of self-perception is often accompanied by depersonalization, that is, the perception of the body as an external, alien. Perhaps an increase in bodily identity and a decrease in the symptom of depersonalization may be caused by self-perception, for example, in a mirror or attention to parts of your body. This requires further research.

This study has limitations. First, the subjects’ attention during the study was not controlled. Using a motor response would provide additional information. Secondly, stimuli were presented in the nested design of the study, which increased the individualization of effects. This can be considered both a disadvantage and a merit of research. Thirdly, the calculation of the localization of distributed activity sources is more accurate with a larger number of electrodes. Fourth, the significance of the face can also be controlled by recording autonomous reactivity, for example, a galvanic skin reaction or by changing the heart rate. Fifth, the study did not control the proximity and duration of friendships.

### Conclusion

In this study, we studied the relationship between brain reactions in the perception of the subject’s own face and the accuracy of perception of interception. Two main results were found. Firstly, the late component of the evoked potential in response to the presentation of the subject’s own face is positively associated with the accuracy of perception of heartbeats, and secondly, the localization of this connection is localized in the right front upper temporal cortex.

